# Site-specific length-biomass relationships of arctic arthropod families are critical for accurate ecological inferences

**DOI:** 10.1101/2023.04.04.534924

**Authors:** Tom S.L. Versluijs, Mikhail K. Zhemchuzhnikov, Dmitry Kutcherov, Tomas Roslin, Niels Martin Schmidt, Jan. A. van Gils, Jeroen Reneerkens

## Abstract

Arthropods play an essential role in terrestrial ecosystems, not least by forming the food base for insectivorous birds. To better understand such trophic interactions, it is essential to monitor seasonal trajectories in arthropod biomass. Because obtaining direct measurements of the body mass of individual specimens is laborious, these data are often indirectly acquired by utilizing allometric length-biomass relationships based on a correlative parameter, such as body length. Studies on insectivorous birds have often used such relationships with a low taxonomic resolution and/or small sample size and/or adopted regressions calibrated in different biomes. Despite the scientific interest in the ecology of arctic arthropods, no site-specific family-level length-biomass relationships have hitherto been published. Here we present 27 family-specific length-biomass relationships from two sites in the High Arctic: Zackenberg in northeast Greenland and Knipovich in north Taimyr, Russia. We show that length-biomass regressions from different sites within the same biome did not affect estimates of phenology but did result in substantially different estimates of arthropod biomass. Estimates of daily biomass at Zackenberg were on average 24% higher when calculated using regressions for Knipovich compared to using regressions for Zackenberg. Our results illustrate that the use of allometric relationships from different sites can significantly alter the biological interpretation of, for instance, the interaction between insectivorous birds and their arthropod prey. We conclude that length-biomass relationships should be locally established rather than being based on global relationships.

**Subjects:** Ecology, Entomology, Terrestrial ecology

## Introduction

Arthropods constitute the most numerically abundant and diverse animal group in terrestrial ecosystems (Goulson, 2019). Across the globe, they play essential roles in nutrient cycling (Yang & Gratton, 2014), and in food webs, where they serve as e.g. pollinators and/or as both prey and predators (Ollerton, Winfree & Tarrant, 2011; Schmidt et al., 2017). For example, ca. 60% of all bird species are insectivorous and rely on arthropods as a resource for survival, growth and egg production (Morse, 1971; Klaassen et al., 2001; Piersma et al., 2003).

Due to the central role of arthropods, long-term ecological monitoring of trends in arthropod diversity and abundance is essential (Hallmann et al., 2017; Goulson, 2019; Gillespie et al., 2020). Detailed information on arthropod abundance is, for instance, key to understanding whether and when the temporal asynchrony between the breeding phenology of insectivorous birds and arthropod availability translates into fitness consequences (Durant et al., 2005; Ramakers, Gienapp & Visser, 2019). Importantly, arthropod taxa, and specimens within taxa, can vary remarkably in their size and morphology (e.g. Baumgärtner & Rothhaupt, 2003; Méthot et al., 2012). Such variation can lead to a lack of correspondence between arthropod abundance and biomass (Saint-Germain et al., 2007). As the quantity of food for insectivorous birds is related to biomass rather than abundance, the monitoring of trends in arthropod biomass will generally allow for more ecologically relevant inferences than monitoring of trends in the mere abundance of arthropods (Saint-Germain et al., 2007; Hallmann et al., 2017).

Acquiring body mass measures for each individual arthropod specimen is, however, laborious. A less time-consuming alternative is to derive estimates of body mass from a correlative parameter such as body length (e.g. Rogers et al. 1977, Sample et al. 1993). This requires knowledge of the allometric relationship between body length and body mass for individual prey taxa. Such allometric relationships are frequently used to provide estimates of prey biomass for insectivorous birds (e.g. Schekkerman et al. 2003, McKinnon et al. 2012, Senner et al. 2017).

Despite the prevalent use of allometric relationships, they have several limitations. In particular, four types of extrapolations can reflect into biased inferences regarding the local availability of arthropod prey to insectivorous birds: (I) Empirically quantified allometric relationships are generally restricted to temperate regions (e.g. Rogers et al. 1977, Schoener 1980, Sample et al. 1993, Sabo et al. 2002), the subtropics (Sage, 1982) or the tropics (Schoener, 1980; Ganihar, 1997; Gruner, 2003), while detailed regressions for other regions – such as the Arctic – are lacking. As arthropods may have specific morphological adaptations to their local environment (e.g. Strathdee & Bale, 1998), applying allometric relations parameterized for one region to another may result in biased estimates of arthropod biomass (Schoener 1980, Hodar 1996, Baumgärtner and Rothhaupt 2003, but see Gowing & Recher, 1984). (II) Empirically quantified allometric relationships are seldom available at a family level or lower taxonomical levels (but see e.g., Sample et al. 1993). As a result, order-level taxonomical equations are frequently used to estimate biomass (e.g. Senner et al. 2017). Resorting to such coarse taxonomic resolution may be problematic, because length-biomass relationships can vary remarkably even within the lower taxonomical levels (Johnston & Cunjak, 1999; Baumgärtner & Rothhaupt, 2003). (III) Empirically quantified allometric relationships are generally based on datasets with limited sample sizes (e.g. Hodar 1996, Sabo et al. 2002). (IV) Empirically quantified allometric relationships are often based on data from several decades ago (e.g. Rogers, Buschbom, and Watson 1977), while the morphology of arthropods may have changed over time (Bowden et al., 2015; Polidori et al., 2020; Wonglersak et al., 2021).

In this study, we present allometric length-biomass relationships at high (family level) taxonomic resolution from two sites in the High Arctic. Drawing on these detailed, site-specific data, we show that estimates of daily arthropod biomass can differ substantially when calculated using length-biomass relationships parameterized for different sites within the same biome. Our results demonstrate the importance of using site-specific length-biomass relationships to enhance the accuracy of biological inference in studies of interactions between insectivorous birds and their arthropod prey.

## Materials & Methods

### Allometric length-biomass regressions

#### Data collection and processing

To derive allometric length-biomass relationships for arctic arthropods at high taxonomical resolution (family level), and to compare the generality of such relations between areas, we used data from two high arctic sites with distinctly different compositions of the arthropod community. Arthropods were caught using yellow pitfalls in June – August 2015 in Zackenberg, northeast Greenland (74°28’ N, 20°34’ W, N=3,594) and June – July 2018 in Knipovich, Taimyr, Russia (76°04’ N, 98°32’ E, N=799). Upon collection, specimens were stored in 96% ethanol and later identified to family level, except for Collembola and Acari which were identified to sub-class level. The length of all specimens was measured under a stereomicroscope to the nearest 0.1 mm directly after taking them out of the ethanol preservative. Lengths were measured from the frons to the tip of the abdomen, excluding any appendages such as antennas, proboscis, or ovipositor. Once measured, all specimens were dried for 2-4 days in open air until their biomass remained constant. All specimens were subsequently oven-dried for 20-24 hours at 60 °C, after which they were placed in a desiccator filled with silica gel to prevent increases in biomass due to moisture absorption. The dry mass of all specimens was weighed directly after taking them out of the desiccator on a microscale balance with an accuracy of 0.01 mg. In general, we aimed to determine the dry mass for each individual specimen, but, to reduce the relative effect of measurement error (Mährlein et al., 2016), we grouped specimens that were too small to be weighed individually into several length classes and subsequently calculated an average dry mass per length class.

#### Fitting statistical models

We fitted separate length-biomass regressions for Zackenberg and Knipovich for all taxonomic groups for which at least eight specimens or length-groups were weighed. A separate regression was fitted to biomass data measured at the level of individuals vs length classes. We fitted four linear models per taxonomic group: (I) an intercept-only model: W = B 0, (II) a linear model on untransformed data: W = B0 + B1 ∗ L, (III) a linear model on natural-log transformed data, i.e. an exponential model: ln(W) = B0 + B1 ∗ L, and (IV) a linear model on natural log-log transformed data, i.e. a power model: ln(W) = B0 + B1 ∗ ln (L), where W = dry mass, L = body length, B0 corresponds to the intercept and B1 to the slope of the linear model. We then selected the best model for each taxonomic group based on AIC (Burnham & Anderson, 2002). To quantify uncertainty for the fitted allometric equations, we calculated 95% quantile confidence intervals for model predictions and regression coefficients using non-parametric (case) bootstrapping using 10,000 bootstrap samples (Efron & Tibshirani, 1994; Nakagawa & Cuthill, 2007). We only used bootstrapping for taxa with a sample size of at least 20. We corrected body mass predictions from log-linear models using Duan’s smearing factor (Duan, 1983; Mährlein et al., 2016). For the full derivation of the appropriate model, dealing with outliers and quantification of model uncertainty, see Article S1.

### Estimates of arthropod phenology and daily biomass based on regressions from two different sites

To establish how our perception of seasonal trajectories in arthropod biomass would differ depending on the origin of the length-biomass regressions employed, we derived estimates of (I) average daily arthropod biomass, and (II) the timing of the median date of arthropod biomass, for 24 years of arthropod data at Zackenberg when biomass was inferred using length–biomass regressions for either Zackenberg or Knipovich.

#### Arthropod data

We analyzed 24 years of arthropod data collected at Zackenberg between 1996 and 2019 (Høye & Forchhammer, 2008; Schmidt et al., 2016). Sampling has occurred at near-weekly intervals from the moment of snowmelt until late August or late September (Schmidt et al., 2016). Arthropods were trapped using yellow pitfall traps at six plots with dimensions 10 × 20m^2^ (Schmidt et al., 2016). One plot was not operational between 1999 and 2018 and was therefore excluded from our analysis. To prevent biases due to interannual differences in the duration of the trapping window, we restricted our analysis to a fixed period from day of year 157 (5-6 June) to 238 (25-26 August). All collected specimens were identified at family-level taxonomic resolution, except for Acari and Collembola which were identified to sub-class level. We restricted our analysis to the taxonomic groups: Araneae *Linyphiidae*, Diptera *Chironomidae*, Dip. *Empididae*, Dip. *Muscidae*, Dip. *Mycetophilidae*, Dip. *Sciaridae*, and Hymenoptera *Ichneumonidae*, for which a length-biomass regression was available for both Zackenberg and Knipovich. This subset included 230,591 specimens, corresponding to 68.7% of the total number of specimens for all taxonomic groups. We excluded Collembola as they made a very limited contribution to overall biomass.

#### Estimating arthropod biomass

The selected Zackenberg arthropod data contain counts of specimens per taxonomic group with a timestamp corresponding to the date when a trap is emptied. These counts thus reflect the cumulative number of specimens collected during all the days for which a trap was active. We first translated this into daily counts per taxonomic group by calculating the average number of trapped specimens per taxonomic group for each day a trap was active. To infer seasonal trajectories in biomass from these count data we then allocated a length to each specimen by random sampling from taxon-specific length distributions (for more details see Article S1). Once a length was allocated to each individual, we used our taxon-specific length-biomass regressions to calculate its corresponding biomass. This latter step was carried out twice, utilizing the regressions specific to either Zackenberg or Knipovich. This resulted in two versions of the Zackenberg arthropod pitfall dataset: one in which biomass was calculated using regressions from Zackenberg and one in which this was calculated using regressions from Knipovich. For both datasets, we then calculated the average arthropod biomass per trap per day for each year. In addition, we estimated arthropod phenology for each year by calculating the date when 50% of cumulative biomass was reached (hereafter “median date of arthropod biomass”) using linear interpolation. We obtained 95% quantile confidence intervals for all estimated parameters using non-parametric (case) bootstrapping with 10,000 bootstrap samples (Efron & Tibshirani, 1994; Nakagawa & Cuthill, 2007). All statistical analyses were performed in R-version R-4.1.2 (R Core Team, 2021).

## Results

### Allometric length-biomass regressions

We identified 4,389 arthropod specimens belonging to 42 taxonomic groups (Zackenberg, n=3,590 specimens of 34 taxonomic groups, and Knipovich, n=799 of 19 taxonomic groups). Body length was measured for 4,383 individual specimens, while biomass was measured for 1,573 individual specimens. The remaining 2,785 individuals, for which biomass could not be individually determined, were grouped into length classes for which an average length and biomass was calculated per group. For 27 taxonomic groups, sample size was sufficiently large to construct allometric relationships for one or both sites, resulting in 22 regressions for Zackenberg and 15 regressions for Knipovich (Table 1). For 31 out of 37 allometric relationships the best supported statistical model was a linear model fitted on natural log-log transformed data, i.e., a power model (Table S2). The allometric relationships of the eight arthropod taxa for which data were available for both Zackenberg and Knipovich are shown in Figure 1, while the relationships for arthropod taxa for which data were only available for either site are shown in Figure S1 and Figure S2, respectively. The average calculated smearing factor across all length-biomass regressions was 1.041 [95% CI, 1.027, 1.058]. Since we used a natural-log transformed response variable, body mass predictions on the arithmetic scale would thus underestimate arthropod biomass by 4.1% (and as much as 25.4% for Ichneumonidae) unless this correction was made (Table 1).

**Table 1.** Best supported allometric length-biomass relationships for 27 arthropod taxa for Zackenberg (n=22) and Knipovich (n=15). The column ‘n’ depicts the number of data points on which each allometric model was fitted and, if applicable, depicts in brackets the sample size before averaging within different length classes. ‘Min (mm)’ and ‘Max (mm)’ indicate the minimum and maximum of length ranges of specimens used to fit each regression. ‘SF’ depicts the smearing factor used to correct back-transformed predictions for models with a natural-log transformed response variable. ‘Level’ indicates whether the allometric model was fitted on individual level measurements or average values for different length classes. ‘Location’ indicates the site where the specimens were collected. Case-bootstrapped 95% confidence intervals for all model parameters can be found in Appendix S1: Table S1. *Abbreviations:* Aca, Acari; Ara, Araneae; Clm, Collembola; Col, Coleoptera; Dip, Diptera; Hym, Hymenoptera; Lep, Lepidoptera; ind, individual level weight measurements; avg, averaged weight estimates per length class; W, body mass (mg); L, body length (mm); KNP, Knipovich; ZAC, Zackenberg.

| Taxon | n | Min<br>(mm) | Max<br>(mm) | B0 | B1 | SF | Level | Model | Location |
| --- | --- | --- | --- | --- | --- | --- | --- | --- | --- |
| Aca sp. | 6 (605) | 0.29 | 1.64 | -3.627 | 2.012 | 1.028 | avg | $\ln(W/SF) = B0 + B1 * \ln(L)$ | ZAC |
| Aca sp. | 9 | 2.08 | 3.24 | -3.438 | 3.249 | 1.008 | ind | $\ln(W/SF) = B0 + B1 * \ln(L)$ | ZAC |
| Ara Dictynidae | 8 | 2.06 | 2.50 | -5.903 | 5.646 | 1.052 | ind | $\ln(W/SF) = B0 + B1 * \ln(L)$ | ZAC |
| Ara Linyphiidae | 28 | 2.30 | 3.70 | -2.422 | 1.928 | 1.013 | ind | $\ln(W/SF) = B0 + B1 * \ln(L)$ | KNP |
| Ara Linyphiidae | 25 | 0.69 | 2.62 | -3.556 | 2.938 | 1.035 | ind | $\ln(W/SF) = B0 + B1 * \ln(L)$ | ZAC |
| Ara Lycosidae | 129 | 1.74 | 8.68 | -3.718 | 2.931 | 1.013 | ind | $\ln(W/SF) = B0 + B1 * \ln(L)$ | ZAC |
| Ara Thomisidae | 12 | 2.55 | 5.55 | -3.268 | 2.963 | 1.007 | ind | $\ln(W/SF) = B0 + B1 * \ln(L)$ | ZAC |
| Clm sp. | 9 (209) | 0.65 | 2.56 | -0.015 | 0.026 | NA | avg | $W = B0 + B1 * L$ | KNP |
| Clm sp. | 8 (1002) | 0.20 | 1.71 | -5.129 | 1.196 | 1.008 | avg | $\ln(W/SF) = B0 + B1 * \ln(L)$ | ZAC |
| Col Carabidae | 21 | 6.30 | 8.30 | 5.371 | NA | NA | ind | $W = B0$ | KNP |
| Col Chrysomelidae | 34 | 4.90 | 6.60 | -3.672 | 3.384 | 1.011 | ind | $\ln(W/SF) = B0 + B1 * \ln(L)$ | KNP |
| Col Staphylinidae | 56 | 2.90 | 7.50 | -4.403 | 2.498 | 1.051 | ind | $\ln(W/SF) = B0 + B1 * \ln(L)$ | KNP |
| Dip Anthomyiidae | 30 | 2.74 | 7.41 | -2.648 | 0.510 | 1.039 | ind | $\ln(W/SF) = B0 + B1 * L$ | ZAC |
| Dip Bolitophilidae | 16 | 3.30 | 4.80 | -4.061 | 1.559 | 1.050 | ind | $\ln(W/SF) = B0 + B1 * \ln(L)$ | KNP |
| Dip Ceratopogonidae | 37 (337) | 1.42 | 2.78 | -3.476 | 0.765 | 1.019 | avg | $\ln(W/SF) = B0 + B1 * \ln(L)$ | ZAC |
| Dip Chironomidae | 83 | 1.16 | 6.10 | -4.738 | 2.414 | 1.150 | ind | $\ln(W/SF) = B0 + B1 * \ln(L)$ | KNP |
| Dip Chironomidae | 67 (328) | 1.31 | 2.93 | -0.028 | 0.031 | NA | avg | $W = B0 + B1 * L$ | ZAC |
| Dip Chironomidae | 45 | 2.70 | 7.64 | -6.286 | 3.127 | 1.058 | ind | $\ln(W/SF) = B0 + B1 * \ln(L)$ | ZAC |
| Dip Culicidae | 78 | 3.97 | 6.94 | -3.093 | 1.313 | 1.098 | ind | $\ln(W/SF) = B0 + B1 * \ln(L)$ | ZAC |
| Dip Empididae | 30 | 5.90 | 7.80 | -2.151 | 1.467 | 1.011 | ind | $\ln(W/SF) = B0 + B1 * \ln(L)$ | KNP |
| Dip Empididae | 22 | 3.57 | 8.50 | -4.295 | 2.350 | 1.025 | ind | $\ln(W/SF) = B0 + B1 * \ln(L)$ | ZAC |
| Dip Muscidae | 44 | 4.00 | 7.80 | -4.685 | 2.949 | 1.013 | ind | $\ln(W/SF) = B0 + B1 * \ln(L)$ | KNP |
| Dip Muscidae | 412 | 3.84 | 8.80 | -4.679 | 2.835 | 1.025 | ind | $\ln(W/SF) = B0 + B1 * \ln(L)$ | ZAC |
| Dip Mycetophilidae | 32 | 3.40 | 5.40 | -3.597 | 1.923 | 1.031 | ind | $\ln(W/SF) = B0 + B1 * \ln(L)$ | KNP |
| Dip Mycetophilidae | 21 | 3.72 | 5.45 | -5.411 | 2.701 | 1.022 | ind | $\ln(W/SF) = B0 + B1 * \ln(L)$ | ZAC |
| Dip Phoridae | 14 | 1.71 | 3.28 | -0.116 | 0.091 | NA | ind | $W = B0 + B1 * L$ | ZAC |
| Dip Scathophagidae | 35 | 5.45 | 8.93 | -3.717 | 2.370 | 1.022 | ind | $\ln(W/SF) = B0 + B1 * \ln(L)$ | ZAC |
| Dip Sciaridae | 110 | 1.56 | 3.50 | -3.854 | 0.525 | 1.047 | ind | $\ln(W/SF) = B0 + B1 * L$ | KNP |
| Dip Sciaridae | 48 (273) | 1.62 | 3.75 | -5.164 | 2.384 | 1.030 | avg | $\ln(W/SF) = B0 + B1 * \ln(L)$ | ZAC |
| Dip Syrphidae | 9 | 6.58 | 12.62 | -5.314 | 3.087 | 1.012 | ind | $\ln(W/SF) = B0 + B1 * \ln(L)$ | ZAC |
| Dip Tachinidae | 11 (28) | 10.56 | 12.48 | -3.952 | 2.617 | 1.005 | avg | $\ln(W/SF) = B0 + B1 * \ln(L)$ | ZAC |
| Dip Tipulidae | 52 | 9.70 | 15.80 | -3.033 | 1.945 | 1.024 | ind | $\ln(W/SF) = B0 + B1 * \ln(L)$ | KNP |
| Dip Trichoceridae | 31 | 3.10 | 5.90 | -6.197 | 3.234 | 1.035 | ind | $\ln(W/SF) = B0 + B1 * \ln(L)$ | KNP |
| Hym Ichneumonidae | 22 | 2.40 | 6.70 | -5.739 | 3.226 | 1.063 | ind | $\ln(W/SF) = B0 + B1 * \ln(L)$ | KNP |
| Hym Ichneumonidae | 50 | 1.86 | 12.34 | -5.559 | 2.928 | 1.254 | ind | $\ln(W/SF) = B0 + B1 * \ln(L)$ | ZAC |
| Hym Tenthredinidae | 12 | 4.30 | 8.10 | -5.434 | 3.261 | 1.061 | ind | $\ln(W/SF) = B0 + B1 * \ln(L)$ | KNP |
| Lep Nymphalidae | 61 | 12.34 | 15.09 | 3.408 | -0.330 | 1.023 | ind | $\ln(W/SF) = B0 + B1 * \ln(L)$ | ZAC |

**Figure 1:**
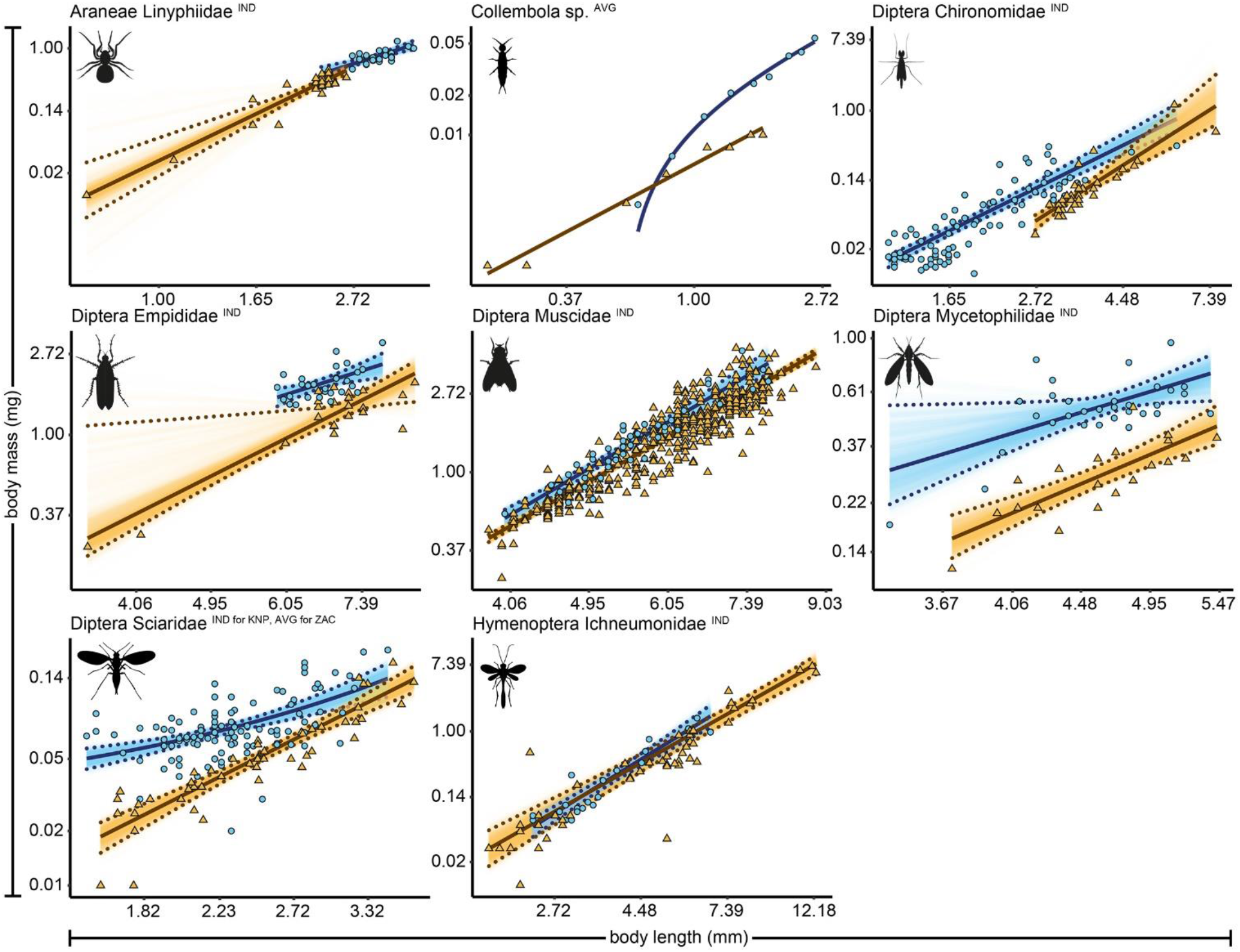
Length-biomass allometric relationship for 8 arthropod taxa for which data were available for both Zackenberg and Knipovich. Data for Zackenberg (ZAC) are depicted as orange triangles and data for Knipovich (KNP) as blue circles. Axes are log transformed but labelled with non-transformed values. Superscripts following the taxonomic names indicate whether datapoints represent individual level weight measurements (‘IND’) or averages per length class (‘AVG’). Solid lines indicate the best supported model for each taxon. Dotted lines indicate 95% quantile confidence intervals calculated over 10,000 case bootstrapping runs. Individual bootstrapping runs are drawn as transparent lines to create a color gradient that visualizes the distribution of best fitting models over all bootstrapping runs. Wide confidence intervals (e.g. for *Empididae*) are an artefact of the use of case-resampling in combination with influential datapoints.

### Estimates of arthropod phenology and abundance based on regressions from two different Sites

Estimates of the average arthropod biomass per trap per day at Zackenberg were on average 23.9% [95% CI: 23.5, 24.4] higher when calculated using regressions for Knipovich than when calculated using regressions for Zackenberg (Fig. 2). When arthropod biomass was calculated using length-biomass regressions for Zackenberg, the median date of arthropod biomass occurred on average 0.13 days [95% CI: 0.03, 0.26] earlier than when regressions for Knipovich were used.

**Figure 2:**
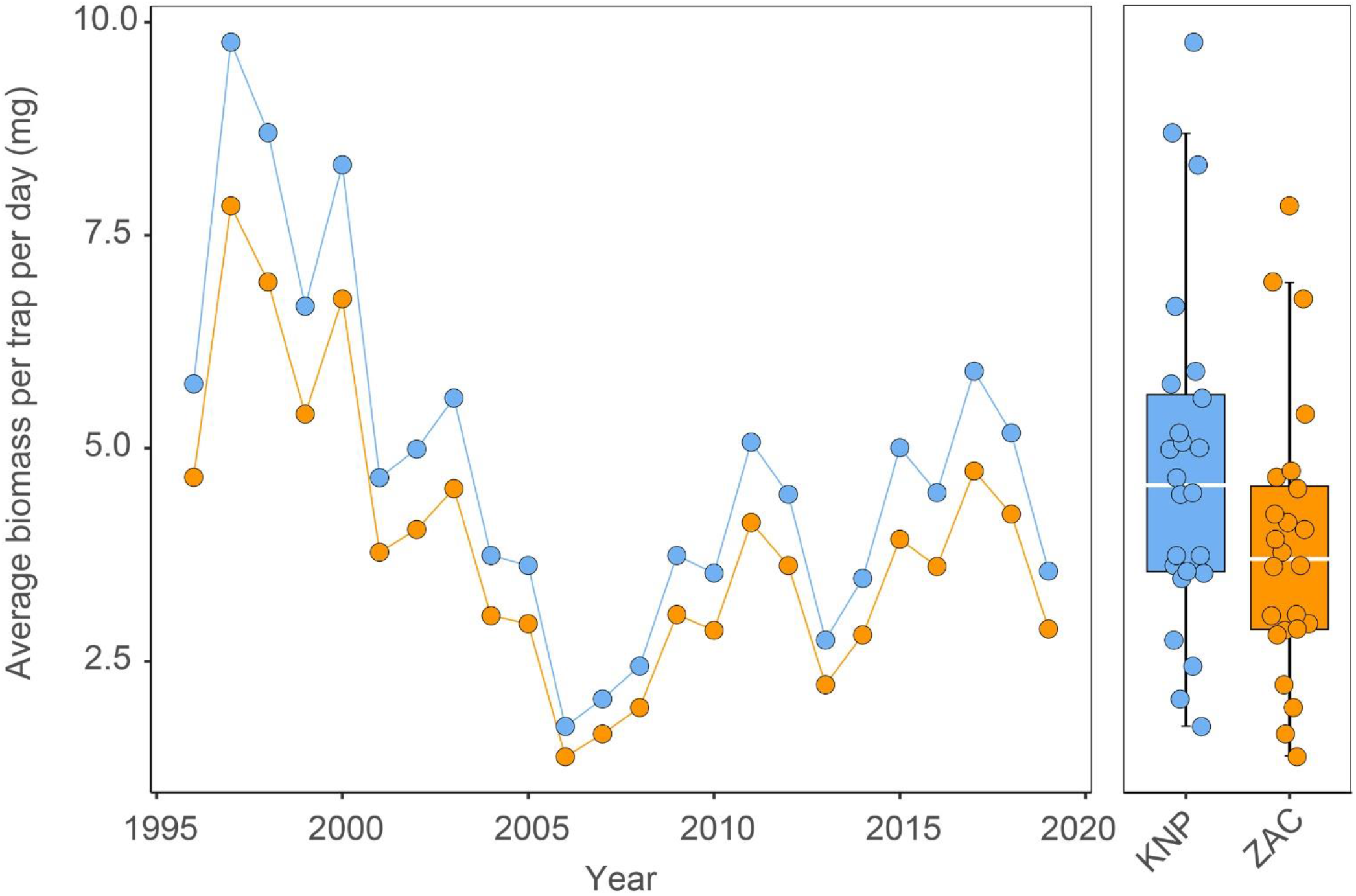
Estimates of average biomass per pitfall trap per day for 24 years of data from Zackenberg (1996 – 2019) when calculated based on regressions for Zackenberg and Knipovich. Data include the Diptera families *Chironomidae, Empididae, Muscidae, Mycetophilidae* and *Sciaridae*, the Hymenoptera family *Ichneumonidae* and the Araneae family *Linyphiidae*. Data depicted in blue are calculated using length-biomass regressions for Knipovich (KNP) and data in orange using regressions for Zackenberg (ZAC). Boxplots summarize the spread in the data, where horizontal white bars indicate the median, the box depicts the interquartile range and whiskers represent 1.5 times the interquartile range from the upper/lower quartile. For visual clarity we applied a horizontal jitter to the raw data depicted in the boxplots.

## Discussion

Based on 24 years of data, we show how the use of allometric relationships from another site within even the same (arctic) biome can result in substantially different estimates of daily arthropod biomass, despite employing identical methodology and taxonomic resolution. This corroborates the findings of earlier studies showing distinct variation within taxonomic groups in regression coefficients or in estimated biomass of invertebrates among sites and/or habitats (e.g. Schoener 1980, Hodar 1996, Baumgärtner and Rothhaupt 2003, but see Gowing and Recher 1984).

Inconsistencies in family-level regression coefficients across sites may arise from differences in site-specific species compositions, or because variation in the time and location of sampling may yield different subsets of sampled species when species differ in their phenology and/or small-scale spatial distribution (Høye & Forchhammer, 2008). In addition, variation in regression parameters might occur due to differences in the timing of emergence among dimorphic sexes (Danks & Oliver, 1972; McLachlan, 1986), or because species differ in their morphological adaptations to their local environment (Strathdee & Bale, 1998). Differences in habitat characteristics and/or food availability may also affect regression parameters by causing intraspecific variation in growth rate (Griffith, Perry & Perry, 1993; Johnston & Cunjak, 1999). Although we employed identical methodologies for both sites, comparisons of regression parameters among studies in general might be hampered by differences in methodologies used for the measuring and weighing of arthropods (Schoener, 1980; Johnston & Cunjak, 1999; Méthot et al., 2012). For instance, corrections for back-transformations from the logarithmic to the arithmetic scale are frequently overlooked (e.g. Sample et al. 1993, Gruner 2003). Variation in body width of specimens might also explain some of the variation between regression parameters, although this may only lead to marginal improvements for allometric relationships constructed at the family level (e.g. Sample et al. 1993, Gruner 2003).

Our results highlight that applying length-biomass relationships calibrated for one site to another site could result in significantly biased estimates of arthropod biomass. Accurate estimates of arthropod biomass are essential to understanding food web dynamics and processes driving community structure (e.g. Saint-Germain et al. 2007) and are for instance crucial in calculating the minimum amount of arthropod biomass required to sustain average growth and survival of shorebird chicks (Schekkerman et al., 2003; Saalfeld et al., 2019). The latter provides a quantitative approach to understand whether asynchrony between insectivorous birds and arthropods translates into fitness consequences for offspring (Durant et al., 2005; Reneerkens et al., 2016). The use of length-biomass regressions from different sites can also affect estimates of the relative contribution of different prey taxa to total prey biomass (Hodar 1996), impacting estimates of prey availability for insectivores. Because our length-biomass equations differ considerably from those from non-arctic regions (e.g. Rogers et al. 1977, Schoener 1980, Sample et al. 1993), we argue that site-specific equations with high taxonomical resolution will provide the most accurate description of local trends in arthropod biomass and will lead to the most accurate biological inference.

Although knowledge on prey biomass can enhance our mechanistic understanding of interactions between insectivores and their prey, the next few steps essential to translating the available prey biomass measured in the field to estimates of the prey biomass actually ingested by consumers is still lacking (Zhemchuzhnikov et al., 2021). First, we should fathom how the number of arthropods recorded relates to what is actually available in the ecosystem, i.e. how observed counts vary with the abundance and activity of arthropods, and how this relationship depends on trapping methods. Second, we should grasp how prey abundance at trapping locations relates to prey abundance at locations where consumers forage. This would call for a mechanistic understanding of the interplay between the movement of the consumer and the spatiotemporal distribution of its resource. Third, we should comprehend how prey abundance at locations where consumers forage translates to ingested prey biomass, i.e. how the diet of the consumers relates to the prey items available (Zhemchuzhnikov et al., 2022).

## Conclusions

We hypothesized that estimates of arthropod biomass in the Arctic were biased by the use of old allometric relationships from other regions and/or by low taxonomical resolution. While the use of allometric relationships from different sites – even within the same biome – had limited effect on estimates of arthropod phenology, they did drastically affect estimates of the locally-available arthropod biomass. As such, this can affect biological interpretations regarding ecological relationships, such as the balance between trophic layers and the food available for offspring growth. Ideally, future studies should establish arthropod length-biomass relationships based on local samples and with high taxonomical resolution.

## Acknowledgements

We thank Aarhus University for providing logistics at Zackenberg. We are grateful for the assistance of many co-workers in the field, and in particular Jannik Hansen and Lars H. Hansen (Zackenberg). Zdenek Gavor and Elin Jørgensen, Aarhus University, sorted and identified arthropods collected in Zackenberg. Arthropods collected in Knipovich were measured and weighed at the Chromas core facility, St. Petersburg State University Research Park. Data from the Greenland Ecosystem Monitoring Programme were provided by the Department of Ecoscience, Aarhus University, Denmark in collaboration with Greenland Institute of Natural Resources, Nuuk, Greenland, and Department of Biology, University of Copenhagen, Denmark.

## Author Contributions

- Tom S. L. Versluijs conceived and designed the research, collected the samples, performed the lab analysis, analysed the data, prepared the figures and/or tables, authored or reviewed drafts of the article, and approved the final draft.
- Mikhail K. Zhemchuzhnikov conceived and designed the research, collected the samples, performed the lab analysis, prepared the figures and/or tables, authored or reviewed drafts of the article, and approved the final draft.
- Dmitry Kutcherov collected the samples, performed the lab analysis, authored or reviewed drafts of the article, and approved the final draft.
- Tomas Roslin authored or reviewed drafts of the article, and approved the final draft.
- Niels Martin Schmidt conceived and designed the research, collected the samples, authored or reviewed drafts of the article, and approved the final draft.
- Jan. A. van Gils authored or reviewed drafts of the article, and approved the final draft.
- Jeroen Reneerkens conceived and designed the research, authored or reviewed drafts of the article, and approved the final draft.

## Data Availability

The following information was supplied regarding data availability: Part of the data utilized for this research is publicly available (Greenland Ecosystem Monitoring, 2020). All other data and/or R-code are provided in a permanent Zenodo repository which can be accessed via the following link: [https://doi.org/10.5281/zenodo.7779505].

