## Supplemental Methods, Tables and Figures for "Site-specific length-biomass relationships of arctic arthropod families are critical for accurate ecological inferences"

### Allometric length-biomass relationships

Dealing with outliers: Before fitting allometric length-biomass regressions we inspected the data for outliers by plotting natural-log transformed biomass against body length. Based on that, we excluded one outlier caused by a measurement error. In addition, we removed 17 datapoints where biomass was not available because of the lower limit of the scales (Mährlein et al. 2016).

Model selection: To be able to compare AIC between models with and without a log-transformed response variable we transformed the AIC value of models with a log-transformed response variable to the arithmetic scale by adding minus twice the logarithm of the Jacobian to the AIC score (Akaike 1978, Burnham and Anderson 2004, Pélabon et al. 2018). Hierarchically more complex models (i.e., nested models with additional parameters) within $\Delta$AIC = 2 of the top-supported model were not considered informative/competitive (Arnold 2010). If multiple models were equally competitive, we preferred the power-model based on residual patterns and biological theoretic foundations.

Correcting back-transformed data: Fitting linear models using least-squares on log-transformed data ensures minimisation of the sum of the squared residuals on the log-transformed scale (Schoener 1980, Mährlein et al. 2016). This implies that body mass estimates from such models correspond to the geometric mean instead of the arithmetic mean, resulting in a bias of the estimates towards smaller values (Hayes and Shonkwiler 2006, Mährlein et al. 2016). To account for this bias, we corrected body mass predictions from log-linear models using Duan's smearing factor (Duan 1983). This factor adjusts the estimated body mass on the arithmetic scale by multiplying it with the mean of the back-transformed residuals of the fitted model (Hayes and Shonkwiler 2006, Mährlein et al. 2016). The corrected equation for the power model then becomes $\ln\left(\frac{W}{SF}\right) = \ln(b0) + b1 * \ln(L)$, where the smearing factor is calculated as $SF = \frac{1}{n}\sum_{i}^{n} e^{\varepsilon_i}$, where $\varepsilon_i$ correspond to the residuals of the fitted model (Mährlein et al. 2016). We refrained from calculating smearing factors when homoscedasticity assumptions were violated. We did not correct for additional biases due to changes in biomass or length as a consequence of preservation in ethanol (Johnston and Cunjak 1999, Mährlein et al. 2016, but see Méthot et al. 2012).

Quantifying model uncertainty: To quantify uncertainty for the fitted allometric equations we calculated 95% quantile confidence intervals for model predictions and regression coefficients using non-parametric (case) bootstrapping using 10,000 bootstrap samples (Efron and Tibshirani 1994, Nakagawa and Cuthill 2007). We opted for non-parametric bootstrapping because normality and homoscedasticity assumptions were violated for several taxa. Although bias-corrected and accelerated bootstrap (BCa) confidence intervals are generally preferred (Efron and Narasimhan 2020), we calculated 95% quantile confidence intervals as the latter resulted in more robust results when dealing with data containing influential datapoints. We only employed bootstrapping for taxa with a sample size of at least n=20.

Calculating biomass: Because two length-biomass regressions are available at Zackenberg for *Chironomidae* (Fig. 1, Table 1), we selected - per specimen - the regression for which the min-max length range of the regression was overlapping with, or was closest to, the length of the specimen.

### Assigning body lengths

The Zackenberg pitfall dataset comprises counts of specimens per taxonomic order, but does not contain measurements on their length or biomass. Thus, in order to use our allometric length-biomass regressions to transform this dataset from counts to biomass, we first needed to assign body lengths to all specimens. This was done by randomly sampling from taxon-specific length distributions in an additional arthropod dataset (n=42,295 specimens of 54 taxonomic groups) that was collected using pitfall traps and sticky traps at Zackenberg in 2015, 2019 and 2021 and in Knipovich in 2018. All specimens in this dataset were identified to family level taxonomic resolution, except for Acari and Collembola which were identified to sub-class taxonomic level. Body lengths of all specimens in this dataset were measured under a stereomicroscope to the nearest 0.1 mm from the frons to the tip of the abdomen (excluding any appendages).

### Supporting Information: Tables

**Table S1: Best supported allometric length-biomass relationships for 27 arthropod taxa including 95% confidence intervals for model parameters.** The column 'n' depicts the number of data points on which each allometric model was fitted and, if applicable, depicts in brackets the sample size before averaging within different length classes. The length range of specimens used for fitting each regression is depicted by the columns 'Min (mm) and 'Max (mm). The columns 'B0', and 'B1' represent model parameters and, if applicable, in square brackets include 95% quantile confidence intervals calculated using non-parametric case-bootstrapping. The column 'SF' depicts a smearing factor that was calculated to correct back-transformed predictions for models with a log transformed response variable. The column 'Level' indicates whether the regression was fitted on individual level measurements or average values for different length classes. The column 'Location' indicates the site where arthropod specimens were collected. *Abbreviations:* ind, individual level weight measurements; avg, averaged weight estimates per length class; W, body mass (mg); SF, smearing factor; L, body length (mm); KNP, Knipovich; ZAC, Zackenberg. *Taxonomic level of Acari and Collembola is subclass.

**Table S2: Results of model selection for 37 allometric relationships for 27 arthropod taxa from Zackenberg and Knipovich.** Models marked in bold were selected as best supported model. This selection generally corresponded to the model with the lowest AIC score, but if multiple models were equally competitive (i.e. $\Delta$AIC<2 with the same number of estimated parameters) we preferred the power-model based on residual patterns and biological theoretic foundations. The column 'n' depicts the number of data points on which each allometric model was fitted. The column 'df' indicates the number of estimated parameters in each model. The column 'logLik' depicts the log-likelihood of each model, and the column 'AIC' shows the AIC score for each model. The column 'Level' indicates whether the regression was fitted on individual level measurements or average values for different length classes. The column 'Location' indicates the site where arthropod specimens were collected. *Abbreviations:* ind, individual level weight measurements; avg, averaged weight estimates per length class; W, body mass (mg); SF, smearing factor; L, body length (mm); KNP, Knipovich; ZAC, Zackenberg.

**Table S1.** Best supported allometric length-biomass relationships for 27 arthropod taxa including 95% confidence intervals for model parameters

| Order | Family | n | Min (mm) | Max (mm) | B0 | B1 | SF | Level | Model | Location |
|---|---|---|---|---|---|---|---|---|---|---|
| Acari* | *sp.* | 6 (605) | 0.29 | 1.64 | -3.627 | 2.012 | 1.028 | avg | ln(W/SF) = B0 + B1 * ln(L) | ZAC |
| Acari* | *sp.* | 9 | 2.08 | 3.24 | -3.438 | 3.249 | 1.008 | ind | ln(W/SF) = B0 + B1 * ln(L) | ZAC |
| Araneae | *Dictynidae* | 8 | 2.06 | 2.50 | -5.903 | 5.646 | 1.052 | ind | ln(W/SF) = B0 + B1 * ln(L) | ZAC |
| Araneae | *Linyphiidae* | 28 | 2.30 | 3.70 | -2.422 [-2.968, -1.608] | 1.928 [1.202, 2.431] | 1.013 [1.007, 1.017] | ind | ln(W/SF) = B0 + B1 * ln(L) | KNP |
| Araneae | *Linyphiidae* | 25 | 0.69 | 2.62 | -3.556 [-4.022, -2.867] | 2.938 [2.114, 3.464] | 1.035 [1.015, 1.053] | ind | ln(W/SF) = B0 + B1 * ln(L) | ZAC |
| Araneae | *Lycosidae* | 129 | 1.74 | 8.68 | -3.718 [-4.087, -3.355] | 2.931 [2.742, 3.120] | 1.013 [1.009, 1.017] | ind | ln(W/SF) = B0 + B1 * ln(L) | ZAC |
| Araneae | *Thomisidae* | 12 | 2.55 | 5.55 | -3.268 | 2.963 | 1.007 | ind | ln(W/SF) = B0 + B1 * ln(L) | ZAC |
| Collembola* | *sp.* | 9 (209) | 0.65 | 2.56 | -0.015 | 0.026 | NA | avg | W = B0 + B1 * L | KNP |
| Collembola* | *sp.* | 8 (1002) | 0.20 | 1.71 | -5.129 | 1.196 | 1.008 | avg | ln(W/SF) = B0 + B1 * ln(L) | ZAC |
| Coleoptera | *Carabidae* | 21 | 6.30 | 8.30 | 5.371 [4.949, 5.802] | NA | NA | ind | W = B0 | KNP |
| Coleoptera | *Chrysomelidae* | 34 | 4.90 | 6.60 | -3.672 [-4.622, -2.616] | 3.384 [2.765, 3.939] | 1.011 [1.006, 1.016] | ind | ln(W/SF) = B0 + B1 * ln(L) | KNP |
| Coleoptera | *Staphylinidae* | 56 | 2.90 | 7.50 | -4.403 [-4.942, -3.851] | 2.498 [2.187, 2.813] | 1.051 [1.035, 1.064] | ind | ln(W/SF) = B0 + B1 * ln(L) | KNP |
| Diptera | *Anthomyiidae* | 30 | 2.74 | 7.41 | -2.648 [-3.147, -2.105] | 0.510 [0.405, 0.597] | 1.039 [1.017, 1.062] | ind | ln(W/SF) = B0 + B1 * L | ZAC |
| Diptera | *Bolitophilidae* | 16 | 3.30 | 4.80 | -4.061 | 1.559 | 1.05 | ind | ln(W/SF) = B0 + B1 * ln(L) | KNP |
| Diptera | *Ceratopogonidae* | 37 (337) | 1.42 | 2.78 | -3.476 [-3.822, -3.207] | 0.765 [0.396, 1.213] | 1.019 [1.010, 1.028] | avg | ln(W/SF) = B0 + B1 * ln(L) | ZAC |
| Diptera | *Chironomidae* | 83 | 1.16 | 6.10 | -4.738 [-4.958, -4.556] | 2.414 [2.171, 2.744] | 1.150 [1.103, 1.192] | ind | ln(W/SF) = B0 + B1 * ln(L) | KNP |
| Diptera | *Chironomidae* | 67 (328) | 1.31 | 2.93 | -0.028 [-0.037, -0.018] | 0.031 [0.027, 0.036] | NA | avg | W = B0 + B1 * L | ZAC |
| Diptera | *Chironomidae* | 45 | 2.70 | 7.64 | -6.286 [-7.578, -5.515] | 3.127 [2.498, 4.199] | 1.058 [1.017, 1.092] | ind | ln(W/SF) = B0 + B1 * ln(L) | ZAC |
| Diptera | *Culicidae* | 78 | 3.97 | 6.94 | -3.093 [-4.184, -2.034] | 1.313 [0.693, 1.958] | 1.098 [1.068, 1.128] | ind | ln(W/SF) = B0 + B1 * ln(L) | ZAC |
| Diptera | *Empididae* | 30 | 5.90 | 7.80 | -2.151 [-3.663, -0.465] | 1.467 [0.574, 2.264] | 1.011 [1.006, 1.016] | ind | ln(W/SF) = B0 + B1 * ln(L) | KNP |
| Diptera | *Empididae* | 22 | 3.57 | 8.50 | -4.295 [-4.995, -0.375] | 2.350 [0.361, 2.741] | 1.025 [1.010, 1.038] | ind | ln(W/SF) = B0 + B1 * ln(L) | ZAC |
| Diptera | *Muscidae* | 44 | 4.00 | 7.80 | -4.685 [-5.406, -4.127] | 2.949 [2.620, 3.373] | 1.013 [1.008, 1.018] | ind | ln(W/SF) = B0 + B1 * ln(L) | KNP |
| Diptera | *Muscidae* | 412 | 3.84 | 8.80 | -4.679 [-4.858, -4.498] | 2.835 [2.733, 2.937] | 1.025 [1.022, 1.029] | ind | ln(W/SF) = B0 + B1 * ln(L) | ZAC |
| Diptera | *Mycetophilidae* | 32 | 3.40 | 5.40 | -3.597 [-5.031, -0.859] | 1.923 [0.168, 2.859] | 1.031 [1.012, 1.046] | ind | ln(W/SF) = B0 + B1 * ln(L) | KNP |
| Diptera | *Mycetophilidae* | 21 | 3.72 | 5.45 | -5.411 [-6.647, -4.085] | 2.701 [1.847, 3.516] | 1.022 [1.009, 1.034] | ind | ln(W/SF) = B0 + B1 * ln(L) | ZAC |
| Diptera | *Phoridae* | 14 | 1.71 | 3.28 | -0.116 | 0.091 | NA | ind | W = B0 + B1 * L | ZAC |
| Diptera | *Scathophagidae* | 35 | 5.45 | 8.93 | -3.717 [-4.641, -2.878] | 2.370 [1.948, 2.839] | 1.022 [1.012, 1.031] | ind | ln(W/SF) = B0 + B1 * ln(L) | ZAC |
| Diptera | *Sciaridae* | 110 | 1.56 | 3.50 | -3.854 [-4.232, -3.452] | 0.525 [0.345, 0.685] | 1.047 [1.033, 1.063] | ind | ln(W/SF) = B0 + B1 * L | KNP |
| Diptera | *Sciaridae* | 48 (273) | 1.62 | 3.75 | -5.164 [-5.570, -4.791] | 2.384 [1.996, 2.799] | 1.030 [1.016, 1.045] | avg | ln(W/SF) = B0 + B1 * ln(L) | ZAC |
| Diptera | *Syrphidae* | 9 | 6.58 | 12.62 | -5.314 | 3.087 | 1.012 | ind | ln(W/SF) = B0 + B1 * ln(L) | ZAC |
| Diptera | *Tachinidae* | 11 (28) | 10.56 | 12.48 | -3.952 | 2.617 | 1.005 | avg | ln(W/SF) = B0 + B1 * ln(L) | ZAC |
| Diptera | *Tipulidae* | 52 | 9.70 | 15.8 | -3.033 [-4.844, -1.373] | 1.945 [1.266, 2.677] | 1.024 [1.012, 1.036] | ind | ln(W/SF) = B0 + B1 * ln(L) | KNP |
| Diptera | *Trichoceridae* | 31 | 3.10 | 5.90 | -6.197 [-7.083, -5.208] | 3.234 [2.490, 3.907] | 1.035 [1.019, 1.047] | ind | ln(W/SF) = B0 + B1 * ln(L) | KNP |
| Hymenoptera | *Ichneumonidae* | 22 | 2.40 | 6.7 | -5.739 [-6.510, -5.132] | 3.226 [2.767, 3.798] | 1.063 [1.024, 1.101] | ind | ln(W/SF) = B0 + B1 * ln(L) | KNP |
| Hymenoptera | *Ichneumonidae* | 50 | 1.86 | 12.34 | -5.559 [-6.170, -4.862] | 2.928 [2.536, 3.273] | 1.254 [1.052, 1.497] | ind | ln(W/SF) = B0 + B1 * ln(L) | ZAC |
| Hymenoptera | *Tenthredinidae* | 12 | 4.30 | 8.1 | -5.434 | 3.261 | 1.061 | ind | ln(W/SF) = B0 + B1 * ln(L) | KNP |
| Lepidoptera | *Nymphalidae* | 61 | 12.34 | 15.09 | 3.408 [0.692, 6.069] | -0.330 [-1.342, 0.702] | 1.023 [1.012, 1.034] | ind | ln(W/SF) = B0 + B1 * ln(L) | ZAC |

**Table S2**. Results of model selection for 37 allometric relationships for 27 arthropod taxa from Zackenberg and Knipovich.

| Order | Family | n | df | logLik | AIC | Level | Model | Location |
|---|---|---|---|---|---|---|---|---|
| **Acari** | **sp.** | **6** | **3** | **0.155** | **-42.746** | **avg** | **ln(Weight/SF) = B0 + B1 * ln(length)** | **ZAC** |
| Acari | sp. | 6 | 3 | 23.151 | -40.302 | avg | Weight = B0 + B1 * length | ZAC |
| Acari | sp. | 6 | 3 | -4.195 | -34.045 | avg | ln(Weight/SF) = B0 + B1 * length | ZAC |
| Acari | sp. | 6 | 2 | 15.369 | -26.737 | avg | Weight = B0 | ZAC |
| Acari | sp. | 9 | 3 | 6.271 | -5.915 | ind | ln(Weight/SF) = B0 + B1 * length | ZAC |
| **Acari** | **sp.** | **9** | **3** | **5.920** | **-5.212** | **ind** | **ln(Weight/SF) = B0 + B1 * ln(length)** | **ZAC** |
| Acari | sp. | 9 | 3 | 4.718 | -3.436 | ind | Weight = B0 + B1 * length | ZAC |
| Acari | sp. | 9 | 2 | -4.737 | 13.475 | ind | Weight = B0 | ZAC |
| Araneae | Dictynidae | 8 | 3 | -2.118 | -10.780 | ind | ln(Weight/SF) = B0 + B1 * length | ZAC |
| **Araneae** | **Dictynidae** | **8** | **3** | **-2.219** | **-10.577** | **ind** | **ln(Weight/SF) = B0 + B1 * ln(length)** | **ZAC** |
| Araneae | Dictynidae | 8 | 3 | 4.203 | -2.406 | ind | Weight = B0 + B1 * length | ZAC |
| Araneae | Dictynidae | 8 | 2 | 1.454 | 1.093 | ind | Weight = B0 | ZAC |
| **Araneae** | **Linyphiidae** | **28** | **3** | **11.905** | **-33.916** | **ind** | **ln(Weight/SF) = B0 + B1 * ln(length)** | **KNP** |
| Araneae | Linyphiidae | 28 | 3 | 11.399 | -32.903 | ind | ln(Weight/SF) = B0 + B1 * length | KNP |
| Araneae | Linyphiidae | 28 | 3 | 17.181 | -28.363 | ind | Weight = B0 + B1 * length | KNP |
| Araneae | Linyphiidae | 28 | 2 | 5.674 | -7.349 | ind | Weight = B0 | KNP |
| **Araneae** | **Linyphiidae** | **25** | **3** | **-1.890** | **-61.322** | **ind** | **ln(Weight/SF) = B0 + B1 * ln(length)** | **ZAC** |
| Araneae | Linyphiidae | 25 | 3 | 32.213 | -58.426 | ind | Weight = B0 + B1 * length | ZAC |
| Araneae | Linyphiidae | 25 | 3 | -6.125 | -52.852 | ind | ln(Weight/SF) = B0 + B1 * length | ZAC |
| Araneae | Linyphiidae | 25 | 2 | 16.380 | -28.760 | ind | Weight = B0 | ZAC |
| **Araneae** | **Lycosidae** | **129** | **3** | **53.263** | **362.152** | **ind** | **ln(Weight/SF) = B0 + B1 * ln(length)** | **ZAC** |
| Araneae | Lycosidae | 129 | 3 | 18.818 | 431.042 | ind | ln(Weight/SF) = B0 + B1 * length | ZAC |
| Araneae | Lycosidae | 129 | 3 | -216.460 | 438.920 | ind | Weight = B0 + B1 * length | ZAC |
| Araneae | Lycosidae | 129 | 2 | -305.091 | 614.181 | ind | Weight = B0 | ZAC |
| Araneae | Thomisidae | 12 | 3 | 0.819 | 4.363 | ind | Weight = B0 + B1 * length | ZAC |
| **Araneae** | **Thomisidae** | **12** | **3** | **8.830** | **5.036** | **ind** | **ln(Weight/SF) = B0 + B1 * ln(length)** | **ZAC** |
| Araneae | Thomisidae | 12 | 3 | 4.433 | 13.830 | ind | ln(Weight/SF) = B0 + B1 * length | ZAC |
| Araneae | Thomisidae | 12 | 2 | -19.365 | 42.730 | ind | Weight = B0 | ZAC |
| **Coleoptera** | **Carabidae** | **21** | **2** | **-29.514** | **63.027** | **ind** | **Weight = B0** | **KNP** |
| Coleoptera | Carabidae | 21 | 3 | -29.488 | 64.975 | ind | Weight = B0 + B1 * length | KNP |
| Coleoptera | Carabidae | 21 | 3 | 5.427 | 65.024 | ind | ln(Weight/SF) = B0 + B1 * ln(length) | KNP |
| Coleoptera | Carabidae | 21 | 3 | 5.381 | 65.116 | ind | ln(Weight/SF) = B0 + B1 * length | KNP |
| **Coleoptera** | **Chrysomelidae** | **34** | **3** | **15.906** | **120.393** | **ind** | **ln(Weight/SF) = B0 + B1 * ln(length)** | **KNP** |
| Coleoptera | Chrysomelidae | 34 | 3 | 15.769 | 120.667 | ind | ln(Weight/SF) = B0 + B1 * length | KNP |
| Coleoptera | Chrysomelidae | 34 | 3 | -59.525 | 125.049 | ind | Weight = B0 + B1 * length | KNP |
| Coleoptera | Chrysomelidae | 34 | 2 | -84.294 | 172.587 | ind | Weight = B0 | KNP |
| **Coleoptera** | **Staphylinidae** | **56** | **3** | **-14.734** | **8.627** | **ind** | **ln(Weight/SF) = B0 + B1 * ln(length)** | **KNP** |
| Coleoptera | Staphylinidae | 56 | 3 | -5.483 | 16.967 | ind | Weight = B0 + B1 * length | KNP |
| Coleoptera | Staphylinidae | 56 | 3 | -20.375 | 19.909 | ind | ln(Weight/SF) = B0 + B1 * length | KNP |
| Coleoptera | Staphylinidae | 56 | 2 | -40.183 | 84.365 | ind | Weight = B0 | KNP |
| **Collembola** | **sp.** | **9** | **3** | **42.927** | **-79.854** | **avg** | **Weight = B0 + B1 * length** | **KNP** |
| Collembola | sp. | 9 | 3 | 3.697 | -72.405 | avg | ln(Weight/SF) = B0 + B1 * ln(length) | KNP |
| Collembola | sp. | 9 | 3 | -2.134 | -60.743 | avg | ln(Weight/SF) = B0 + B1 * length | KNP |
| Collembola | sp. | 9 | 2 | 24.247 | -44.494 | avg | Weight = B0 | KNP |
| **Collembola** | **sp.** | **8** | **3** | **5.226** | **-92.032** | **avg** | **ln(Weight/SF) = B0 + B1 * ln(length)** | **ZAC** |
| Collembola | sp. | 8 | 3 | 48.657 | -91.314 | avg | Weight = B0 + B1 * length | ZAC |
| Collembola | sp. | 8 | 3 | -1.540 | -78.499 | avg | ln(Weight/SF) = B0 + B1 * length | ZAC |
| Collembola | sp. | 8 | 2 | 33.828 | -63.655 | avg | Weight = B0 | ZAC |
| **Diptera** | **Anthomyiidae** | **30** | **3** | **-5.605** | **24.705** | **ind** | **ln(Weight/SF) = B0 + B1 * length** | **ZAC** |
| Diptera | Anthomyiidae | 30 | 3 | -7.167 | 27.828 | ind | ln(Weight/SF) = B0 + B1 * ln(length) | ZAC |
| Diptera | Anthomyiidae | 30 | 3 | -20.179 | 46.359 | ind | Weight = B0 + B1 * length | ZAC |
| Diptera | Anthomyiidae | 30 | 2 | -36.853 | 77.706 | ind | Weight = B0 | ZAC |
| **Diptera** | **Bolitophilidae** | **16** | **3** | **-3.573** | **-45.068** | **ind** | **ln(Weight/SF) = B0 + B1 * ln(length)** | **KNP** |
| Diptera | Bolitophilidae | 16 | 3 | -3.608 | -44.998 | ind | ln(Weight/SF) = B0 + B1 * length | KNP |
| Diptera | Bolitophilidae | 16 | 3 | 21.453 | -36.905 | ind | Weight = B0 + B1 * length | KNP |
| Diptera | Bolitophilidae | 16 | 2 | 20.189 | -36.378 | ind | Weight = B0 | KNP |
| Diptera | Ceratopogonidae | 37 | 3 | 116.997 | -227.994 | avg | Weight = B0 + B1 * length | ZAC |
| Diptera | Ceratopogonidae | 37 | 3 | 7.989 | -227.752 | avg | ln(Weight/SF) = B0 + B1 * length | ZAC |
| **Diptera** | **Ceratopogonidae** | **37** | **3** | **7.471** | **-226.717** | **avg** | **ln(Weight/SF) = B0 + B1 * ln(length)** | **ZAC** |
| Diptera | Ceratopogonidae | 37 | 2 | 109.190 | -214.381 | avg | Weight = B0 | ZAC |
| **Diptera** | **Chironomidae** | **83** | **3** | **-62.704** | **-395.525** | **ind** | **ln(Weight/SF) = B0 + B1 * ln(length)** | **KNP** |
| Diptera | Chironomidae | 83 | 3 | -72.143 | -376.647 | ind | ln(Weight/SF) = B0 + B1 * length | KNP |
| Diptera | Chironomidae | 83 | 3 | 134.368 | -262.737 | ind | Weight = B0 + B1 * length | KNP |
| Diptera | Chironomidae | 83 | 2 | 89.769 | -175.537 | ind | Weight = B0 | KNP |
| **Diptera** | **Chironomidae** | **67** | **3** | **217.704** | **-429.408** | **avg** | **Weight = B0 + B1 * length** | **ZAC** |
| Diptera | Chironomidae | 67 | 3 | -15.192 | -418.686 | avg | ln(Weight/SF) = B0 + B1 * ln(length) | ZAC |
| Diptera | Chironomidae | 67 | 3 | -17.342 | -414.387 | avg | ln(Weight/SF) = B0 + B1 * length | ZAC |
| Diptera | Chironomidae | 67 | 2 | 176.776 | -349.551 | avg | Weight = B0 | ZAC |

| | | | | | | | | |
|---|---|---|---|---|---|---|---|---|
| **Diptera** | **Chironomidae** | **45** | **3** | **-11.224** | **-183.478** | **ind** | **ln(Weight/SF) = B0 + B1 * ln(length)** | **ZAC** |
| Diptera | Chironomidae | 45 | 3 | -16.205 | -173.514 | ind | ln(Weight/SF) = B0 + B1 * length | ZAC |
| Diptera | Chironomidae | 45 | 3 | 33.810 | -61.621 | ind | Weight = B0 + B1 * length | ZAC |
| Diptera | Chironomidae | 45 | 2 | 13.178 | -22.357 | ind | Weight = B0 | ZAC |
| **Diptera** | **Culicidae** | **78** | **3** | **-43.819** | **-46.284** | **ind** | **ln(Weight/SF) = B0 + B1 * ln(length)** | **ZAC** |
| Diptera | Culicidae | 78 | 3 | -44.341 | -45.240 | ind | ln(Weight/SF) = B0 + B1 * length | ZAC |
| Diptera | Culicidae | 78 | 3 | 12.576 | -19.153 | ind | Weight = B0 + B1 * length | ZAC |
| Diptera | Culicidae | 78 | 2 | 9.210 | -14.420 | ind | Weight = B0 | ZAC |
| Diptera | Empididae | 30 | 3 | 14.898 | 14.560 | ind | ln(Weight/SF) = B0 + B1 * length | KNP |
| **Diptera** | **Empididae** | **30** | **3** | **14.623** | **15.111** | **ind** | **ln(Weight/SF) = B0 + B1 * ln(length)** | **KNP** |
| Diptera | Empididae | 30 | 3 | -7.227 | 20.453 | ind | Weight = B0 + B1 * length | KNP |
| Diptera | Empididae | 30 | 2 | -13.577 | 31.153 | ind | Weight = B0 | KNP |
| Diptera | Empididae | 22 | 3 | -1.900 | 9.799 | ind | Weight = B0 + B1 * length | ZAC |
| **Diptera** | **Empididae** | **22** | **3** | **1.732** | **10.367** | **ind** | **ln(Weight/SF) = B0 + B1 * ln(length)** | **ZAC** |
| Diptera | Empididae | 22 | 3 | -1.753 | 17.338 | ind | ln(Weight/SF) = B0 + B1 * length | ZAC |
| Diptera | Empididae | 22 | 2 | -12.229 | 28.457 | ind | Weight = B0 | ZAC |
| **Diptera** | **Muscidae** | **44** | **3** | **17.467** | **3.390** | **ind** | **ln(Weight/SF) = B0 + B1 * ln(length)** | **KNP** |
| Diptera | Muscidae | 44 | 3 | 15.545 | 7.234 | ind | ln(Weight/SF) = B0 + B1 * length | KNP |
| Diptera | Muscidae | 44 | 3 | -16.393 | 38.787 | ind | Weight = B0 + B1 * length | KNP |
| Diptera | Muscidae | 44 | 2 | -55.825 | 115.649 | ind | Weight = B0 | KNP |
| **Diptera** | **Muscidae** | **412** | **3** | **35.168** | **276.004** | **ind** | **ln(Weight/SF) = B0 + B1 * ln(length)** | **ZAC** |
| Diptera | Muscidae | 412 | 3 | 21.845 | 302.651 | ind | ln(Weight/SF) = B0 + B1 * length | ZAC |
| Diptera | Muscidae | 412 | 3 | -267.639 | 541.279 | ind | Weight = B0 + B1 * length | ZAC |
| Diptera | Muscidae | 412 | 2 | -543.813 | 1091.625 | ind | Weight = B0 | ZAC |
| **Diptera** | **Mycetophilidae** | **32** | **3** | **0.154** | **-36.849** | **ind** | **ln(Weight/SF) = B0 + B1 * ln(length)** | **KNP** |
| Diptera | Mycetophilidae | 32 | 3 | -0.669 | -35.203 | ind | ln(Weight/SF) = B0 + B1 * length | KNP |
| Diptera | Mycetophilidae | 32 | 3 | 20.313 | -34.626 | ind | Weight = B0 + B1 * length | KNP |
| Diptera | Mycetophilidae | 32 | 2 | 16.138 | -28.275 | ind | Weight = B0 | KNP |
| **Diptera** | **Mycetophilidae** | **21** | **3** | **3.498** | **-55.602** | **ind** | **ln(Weight/SF) = B0 + B1 * ln(length)** | **ZAC** |
| Diptera | Mycetophilidae | 21 | 3 | 3.157 | -54.921 | ind | ln(Weight/SF) = B0 + B1 * length | ZAC |
| Diptera | Mycetophilidae | 21 | 3 | 28.245 | -50.491 | ind | Weight = B0 + B1 * length | ZAC |
| Diptera | Mycetophilidae | 21 | 2 | 19.980 | -35.960 | ind | Weight = B0 | ZAC |
| **Diptera** | **Phoridae** | **14** | **3** | **43.617** | **-81.233** | **ind** | **Weight = B0 + B1 * length** | **ZAC** |
| Diptera | Phoridae | 14 | 3 | 2.307 | -76.059 | ind | ln(Weight/SF) = B0 + B1 * ln(length) | ZAC |
| Diptera | Phoridae | 14 | 3 | 1.455 | -74.354 | ind | ln(Weight/SF) = B0 + B1 * length | ZAC |
| Diptera | Phoridae | 14 | 2 | 25.754 | -47.508 | ind | Weight = B0 | ZAC |
| Diptera | Scathophagidae | 35 | 3 | 4.987 | 54.201 | ind | ln(Weight/SF) = B0 + B1 * length | ZAC |
| **Diptera** | **Scathophagidae** | **35** | **3** | **4.893** | **54.388** | **ind** | **ln(Weight/SF) = B0 + B1 * ln(length)** | **ZAC** |
| Diptera | Scathophagidae | 35 | 3 | -26.074 | 58.149 | ind | Weight = B0 + B1 * length | ZAC |
| Diptera | Scathophagidae | 35 | 2 | -49.278 | 102.556 | ind | Weight = B0 | ZAC |
| **Diptera** | **Sciaridae** | **110** | **3** | **-28.516** | **-515.981** | **ind** | **ln(Weight/SF) = B0 + B1 * length** | **KNP** |
| Diptera | Sciaridae | 110 | 3 | -30.999 | -511.015 | ind | ln(Weight/SF) = B0 + B1 * ln(length) | KNP |
| Diptera | Sciaridae | 110 | 3 | 248.534 | -491.067 | ind | Weight = B0 + B1 * length | KNP |
| Diptera | Sciaridae | 110 | 2 | 223.283 | -442.566 | ind | Weight = B0 | KNP |
| **Diptera** | **Sciaridae** | **48** | **3** | **-2.006** | **-280.922** | **avg** | **ln(Weight/SF) = B0 + B1 * ln(length)** | **ZAC** |
| Diptera | Sciaridae | 48 | 3 | -2.791 | -279.352 | avg | ln(Weight/SF) = B0 + B1 * length | ZAC |
| Diptera | Sciaridae | 48 | 3 | 129.493 | -252.985 | avg | Weight = B0 + B1 * length | ZAC |
| Diptera | Sciaridae | 48 | 2 | 92.578 | -181.156 | avg | Weight = B0 | ZAC |
| **Diptera** | **Syrphidae** | **9** | **3** | **4.227** | **21.283** | **ind** | **ln(Weight/SF) = B0 + B1 * ln(length)** | **ZAC** |
| Diptera | Syrphidae | 9 | 3 | 3.240 | 23.258 | ind | ln(Weight/SF) = B0 + B1 * length | ZAC |
| Diptera | Syrphidae | 9 | 3 | -11.500 | 29.000 | ind | Weight = B0 + B1 * length | ZAC |
| Diptera | Syrphidae | 9 | 2 | -24.852 | 53.703 | ind | Weight = B0 | ZAC |
| Diptera | Tachinidae | 11 | 3 | 9.526 | 40.882 | avg | ln(Weight/SF) = B0 + B1 * length | ZAC |
| **Diptera** | **Tachinidae** | **11** | **3** | **9.470** | **40.993** | **avg** | **ln(Weight/SF) = B0 + B1 * ln(length)** | **ZAC** |
| Diptera | Tachinidae | 11 | 3 | -18.299 | 42.598 | avg | Weight = B0 + B1 * length | ZAC |
| Diptera | Tachinidae | 11 | 2 | -23.351 | 50.703 | avg | Weight = B0 | ZAC |
| Diptera | Tipulidae | 52 | 3 | 6.949 | 181.089 | ind | ln(Weight/SF) = B0 + B1 * length | KNP |
| **Diptera** | **Tipulidae** | **52** | **3** | **6.859** | **181.270** | **ind** | **ln(Weight/SF) = B0 + B1 * ln(length)** | **KNP** |
| Diptera | Tipulidae | 52 | 3 | -102.319 | 210.637 | ind | Weight = B0 + B1 * length | KNP |
| Diptera | Tipulidae | 52 | 2 | -117.763 | 239.526 | ind | Weight = B0 | KNP |
| **Diptera** | **Trichoceridae** | **31** | **3** | **-1.856** | **-99.172** | **ind** | **ln(Weight/SF) = B0 + B1 * ln(length)** | **KNP** |
| Diptera | Trichoceridae | 31 | 3 | -2.267 | -98.350 | ind | ln(Weight/SF) = B0 + B1 * length | KNP |
| Diptera | Trichoceridae | 31 | 3 | 41.458 | -76.915 | ind | Weight = B0 + B1 * length | KNP |
| Diptera | Trichoceridae | 31 | 2 | 22.682 | -41.364 | ind | Weight = B0 | KNP |
| **Hymenoptera** | **Ichneumonidae** | **22** | **3** | **-7.329** | **-51.531** | **ind** | **ln(Weight/SF) = B0 + B1 * ln(length)** | **KNP** |
| Hymenoptera | Ichneumonidae | 22 | 3 | -8.281 | -49.627 | ind | ln(Weight/SF) = B0 + B1 * length | KNP |
| Hymenoptera | Ichneumonidae | 22 | 3 | 12.081 | -18.162 | ind | Weight = B0 + B1 * length | KNP |
| Hymenoptera | Ichneumonidae | 22 | 2 | -10.494 | 24.987 | ind | Weight = B0 | KNP |
| **Hymenoptera** | **Ichneumonidae** | **50** | **3** | **-46.965** | **-9.878** | **ind** | **ln(Weight/SF) = B0 + B1 * ln(length)** | **ZAC** |

| | | | | | | | | |
|---|---|---|---|---|---|---|---|---|
| Hymenoptera | Ichneumonidae | 50 | 3 | -51.817 | -0.173 | ind | ln(Weight/SF) = B0 + B1 * length | ZAC |
| Hymenoptera | Ichneumonidae | 50 | 3 | -57.169 | 120.338 | ind | Weight = B0 + B1 * length | ZAC |
| Hymenoptera | Ichneumonidae | 50 | 2 | -92.948 | 189.896 | ind | Weight = B0 | ZAC |
| **Hymenoptera** | **Tenthredinidae** | **12** | **3** | **-3.602** | **15.163** | **ind** | **ln(Weight/SF) = B0 + B1 * ln(length)** | **KNP** |
| Hymenoptera | Tenthredinidae | 12 | 3 | -3.657 | 15.274 | ind | ln(Weight/SF) = B0 + B1 * length | KNP |
| Hymenoptera | Tenthredinidae | 12 | 3 | -11.926 | 29.852 | ind | Weight = B0 + B1 * length | KNP |
| Hymenoptera | Tenthredinidae | 12 | 2 | -19.607 | 43.215 | ind | Weight = B0 | KNP |
| **Lepidoptera** | **Nymphalidae** | **61** | **3** | **10.275** | **295.749** | **ind** | **ln(Weight/SF) = B0 + B1 * ln(length)** | **ZAC** |
| Lepidoptera | Nymphalidae | 61 | 3 | 10.275 | 295.749 | ind | ln(Weight/SF) = B0 + B1 * length | ZAC |
| Lepidoptera | Nymphalidae | 61 | 2 | -154.053 | 312.106 | ind | Weight = B0 | ZAC |
| Lepidoptera | Nymphalidae | 61 | 3 | -153.823 | 313.646 | ind | Weight = B0 + B1 * length | ZAC |

### Supporting Information: Figures

**Figure S1: Length-biomass allometric relationship for 13 arthropod taxa for which data were only available for Zackenberg.** Axes are log transformed but labelled with non-transformed values to ease interpretation. Superscripts following taxonomic names indicate whether datapoints represent individual level weight measurements ('IND') or averages per length group ('AVG'). Note that a separate allometric relationship is presented for small Acari for which data are based on averages per length class, and for large Acari for which data are based on individual measurements. Raw datapoints are depicted as orange triangles. Solid lines indicate the best supported model for each taxon. Dotted lines indicate 95% quantile confidence intervals calculated over 10,000 case bootstrapping runs. Individual bootstrapping runs are drawn as transparent lines to create a color gradient that visualizes the distribution of best fitting models over all bootstrapping runs.

**Figure S2: Length-biomass allometric relationship for 7 arthropod taxa for which data were only available for Knipovich**. Axes are log transformed but labelled with non-transformed values to ease interpretation. Superscripts following taxonomic names indicate whether datapoints represent individual level weight measurements ('IND') or averages per length group ('AVG'). Raw datapoints are depicted as blue circles. Solid lines indicate the best supported model for each taxon. Dotted lines indicate 95% quantile confidence intervals calculated over 10,000 case bootstrapping runs. Individual bootstrapping runs are drawn as transparent lines to create a color gradient that visualizes the distribution of best fitting models over all bootstrapping runs.

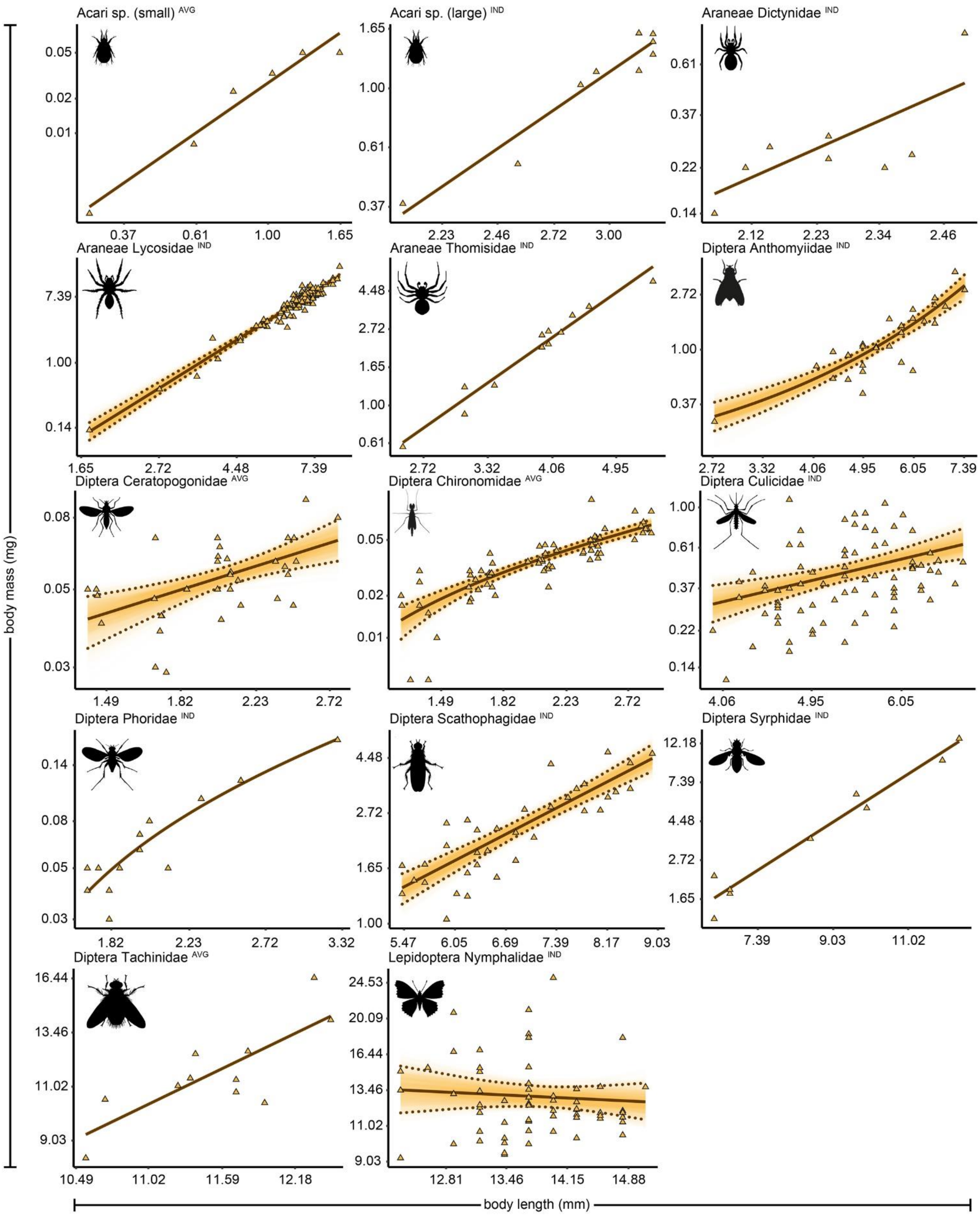
Acari sp. (small) AVG
Acari sp. (large) IND
Araneae Dictynidae IND
Araneae Lycosidae IND
Araneae Thomisidae IND
Diptera Anthomyiidae IND
Diptera Ceratopogonidae AVG
Diptera Chironomidae AVG
Diptera Culicidae IND
Diptera Phoridae IND
Diptera Scathophagidae IND
Diptera Syrphidae IND
Diptera Tachinidae AVG
Lepidoptera Nymphalidae IND
body mass (mg)
body length (mm)

Figure S1

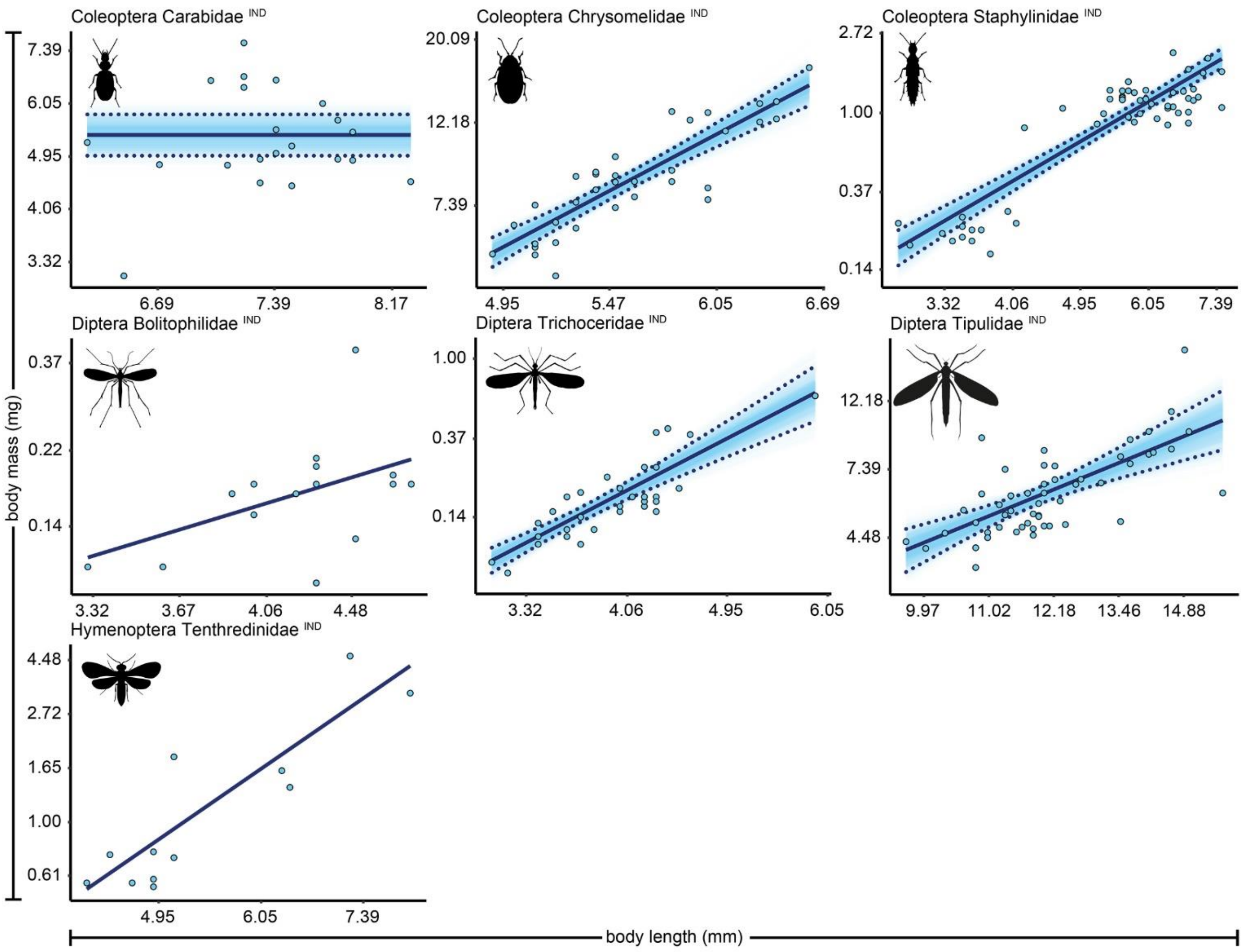

Coleoptera Carabidae IND
Coleoptera Chrysomelidae IND
Coleoptera Staphylinidae IND
Diptera Bolitophilidae IND
Diptera Trichoceridae IND
Diptera Tipulidae IND
Hymenoptera Tenthredinidae IND
body mass (mg)
body length (mm)


Figure S2
